# Dromi: Python package for parallel computation of similarity measures among vector-encoded sequences

**DOI:** 10.1101/2023.07.05.547866

**Authors:** Lys Sanz Moreta

## Abstract

Calculating similarities among sequences (i.e biological sequences) can be a challenging task. Here I introduce Dromi, a simple python package that can compute different similarity measurements (i.e percent identity, cosine similarity, kmer similarities) across aligned vector-encoded sequences. This is a crucial step required to perform both upstream and downstream sequence machine learning tasks such as sequence clustering [1, 2, 3], sequence analysis [4] and other pre- or post-processing demands on sequences. Additionally, this package introduces the calculation of the measure referred as *positional weights*. These represent the cosine similarities or residue-conservation across sequence elements (i.e amino acids in peptide sequences) in the same site (column). The program can also deal with sequences of variable length since end-padded positions are not considered for the calculations. The presented implementations are an incorporation into the arsenal of tools to measure similarity among small peptide sequences such as epitopes.

## 1 Introduction

Measuring the similarity between the sequences is not a trivial task. Currently, most packages solely offer measurements such as percent identity, kmer-based distances (Hobohm algorithm [5]), Levenshtein distances [6] or some more advanced and expensive methodologies used in biology that include biological features [7]. Dromi is a simple python package that can compute the cosine similarity, between vector-encoded sequences in a paralleled and batched manner using standard python libraries.

Calculating sequence similarity is an essential step *a priori* or *posteriori* for performing machine learning tasks such as data partitioning, clustering or sequence pattern discovery. Dromi contains the implementations for classical sequence comparison measurements between aligned sequences such as percent identity. Nonetheless, in addition it offers more exact similarity measurements such as cosine similarity. Moreover, it introduces the novel *positional weights*, meaning the cosine similarities as a measure of conservation [4] across sequence elements such as residues in aligned biological sequences at the same position (see 3.3).

Positional weights are designed to i) become an easy visual assessment of residue conservation ii) an indicator or weight to mask certain positions, for example during a machine learning task. The package also offers additional functions such as biological sequence encoding and sequence padding. The package is expected to expand with new utilities for treatment of sequences in machine learning.

## 2 Notation table

## 3 Available Implementations

The presented python package offers the following computations. Note that the *for* loops are optimized making use of *numpy* broadcasting capabilities and python native functions (such as *map* and *multiprocessing*). This makes it a simple light package with low requirements for installation.

### 3.1 Single element

#### 3.1.1 Percent Identity Mean

The percent identity average value is a numeric score calculated for each pair of aligned sequences. The score represents the number of identical residues (“matches”) in relation to the length of the alignment. The score increases one unit when the compared elements are identical and there is no padded elements (missing elements in the aligned sequence).

##### Algorithm 1 Percent Identity Mean between a pair of sequences. Computational complexity 𝒪(*L*^2^)

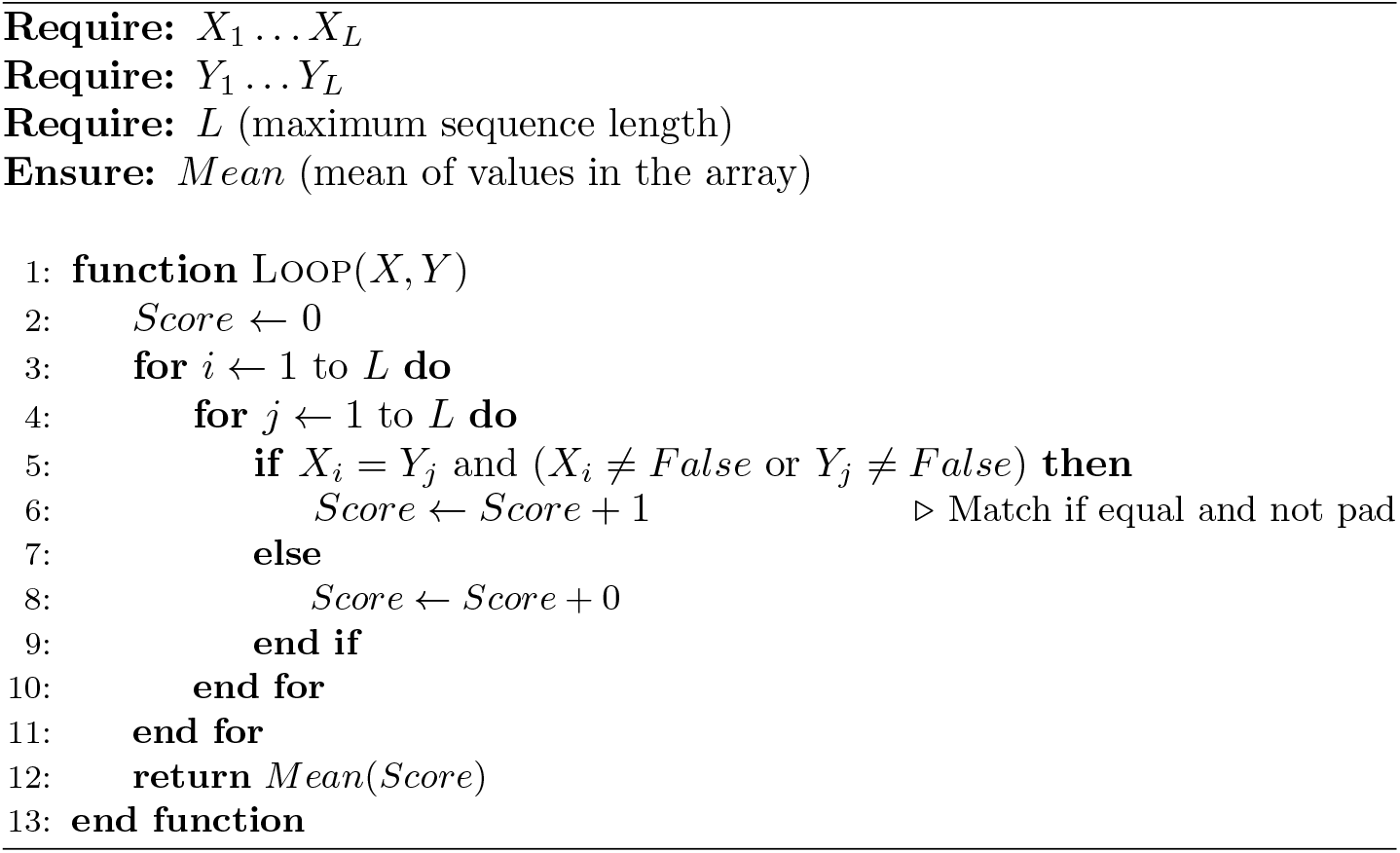

#### 3.1.2 Cosine similarity mean

The cosine similarity is the measurement of similarity among two vectors regardless of their difference in magnitude or direction. Particularly, it measures the similarity in direction or orientation of the compared vectors. The analyzed vectors need to be compared in the same inner product space, meaning that their inner product results into a scalar value. Then, the similarity of the two vectors is measured by the cosine of the angle between them.

The similarity values range from -1 to 1. Similarity scores of 1 indicate higher degree of similarity due to the angle between the vectors being 0. Scores of 0 indicate that the vectors are perpendicular to each other, whilst values of - 1 indicate that the vectors follow opposite directions. The cosine similarity between 2 vectors 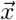 and 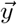 is shown in Equation 1.

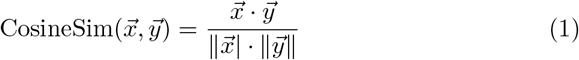

##### Algorithm 2 Cosine Similarity Mean between a pair of sequences 𝒪(*L*^2^)

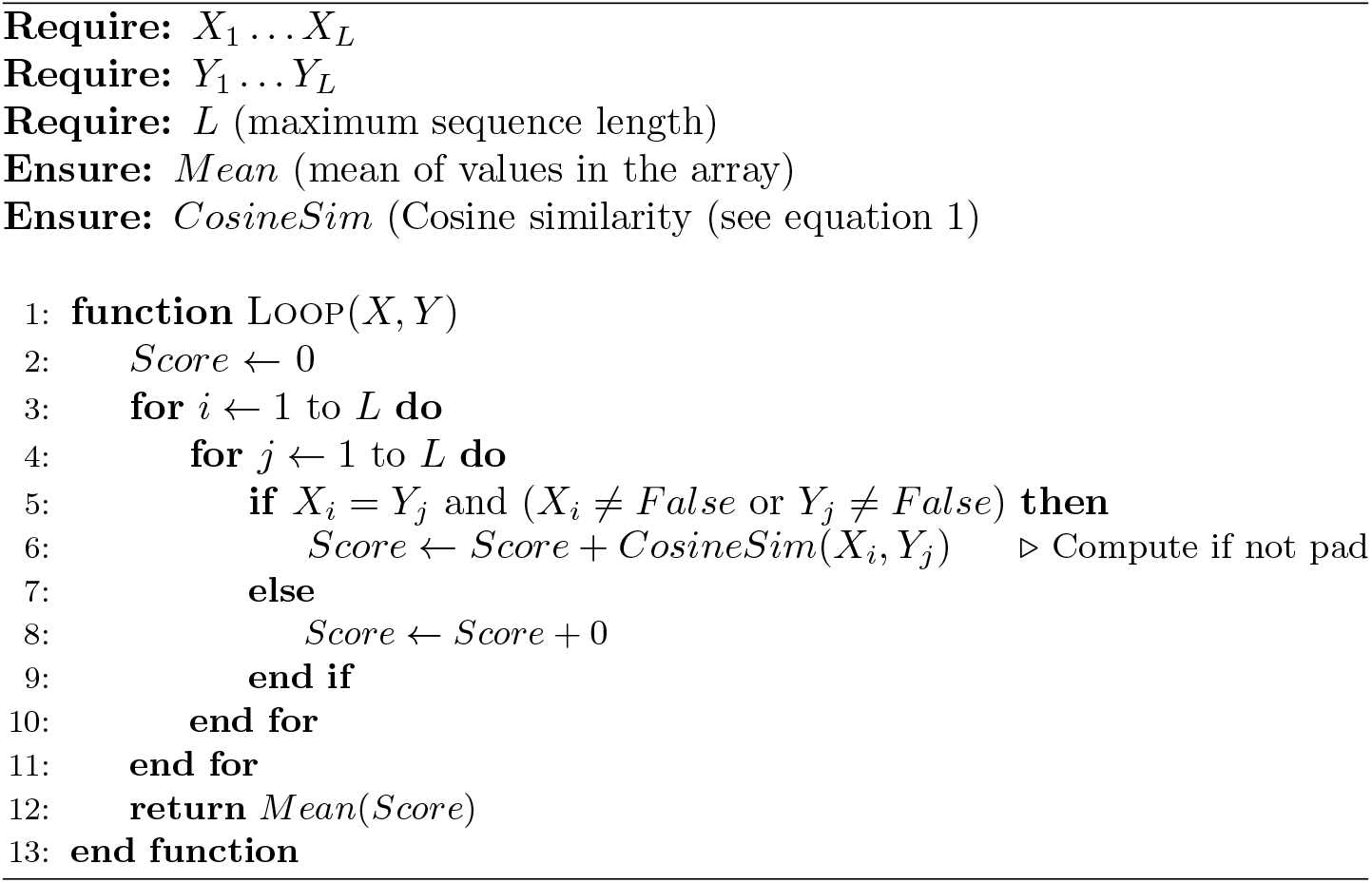

### 3.2 Kmer computations

In the following implementations, the sequences are subdivided onto overlapping fragments referred commonly as *kmers*. This computation allows for finding similarities among neighbour sub-sequences. The larger the *kmer* size the smaller number of computed kmers.

#### 3.2.1 Kmer Percent Identity Mean

This calculation is analogous to the one described in section 3.1.1 however in this occasion the *kmers* of the sequence are compared instead of the entire sequence.

##### Algorithm 3 Kmers Percent Identity Mean between a pair of sequences 𝒪(*NumK*^2^*K*^2^)

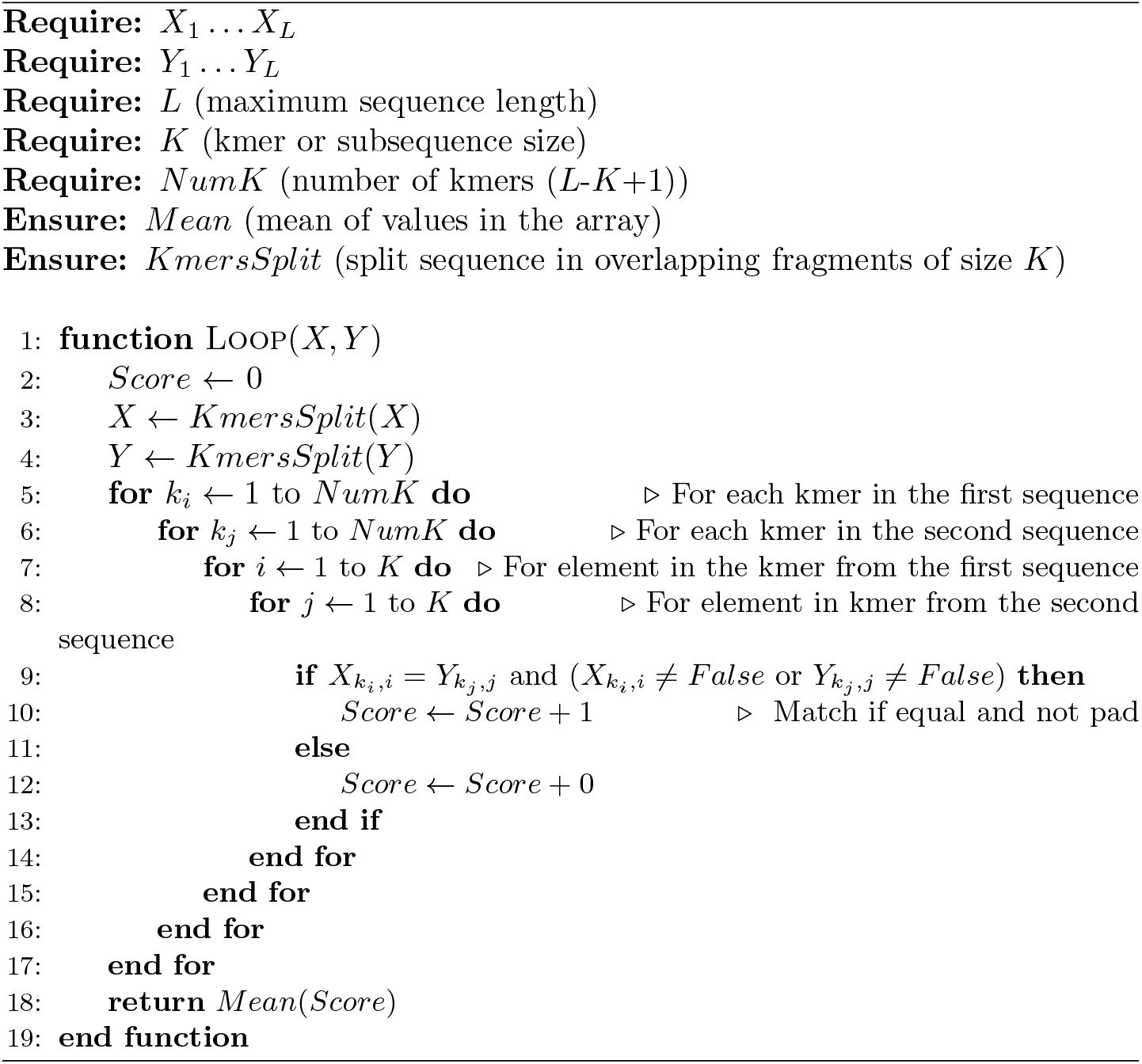

#### 3.2.2 Kmers Cosine Similarity Mean

This calculation is analogous to the one described in section 3.1.2 however in this occasion the *kmers* of the sequence are compared instead of the entire sequence.

##### Algorithm 4 Kmers Cosine similarity Mean between a pair of sequences. 𝒪(*NumK*^2^*K*^2^)

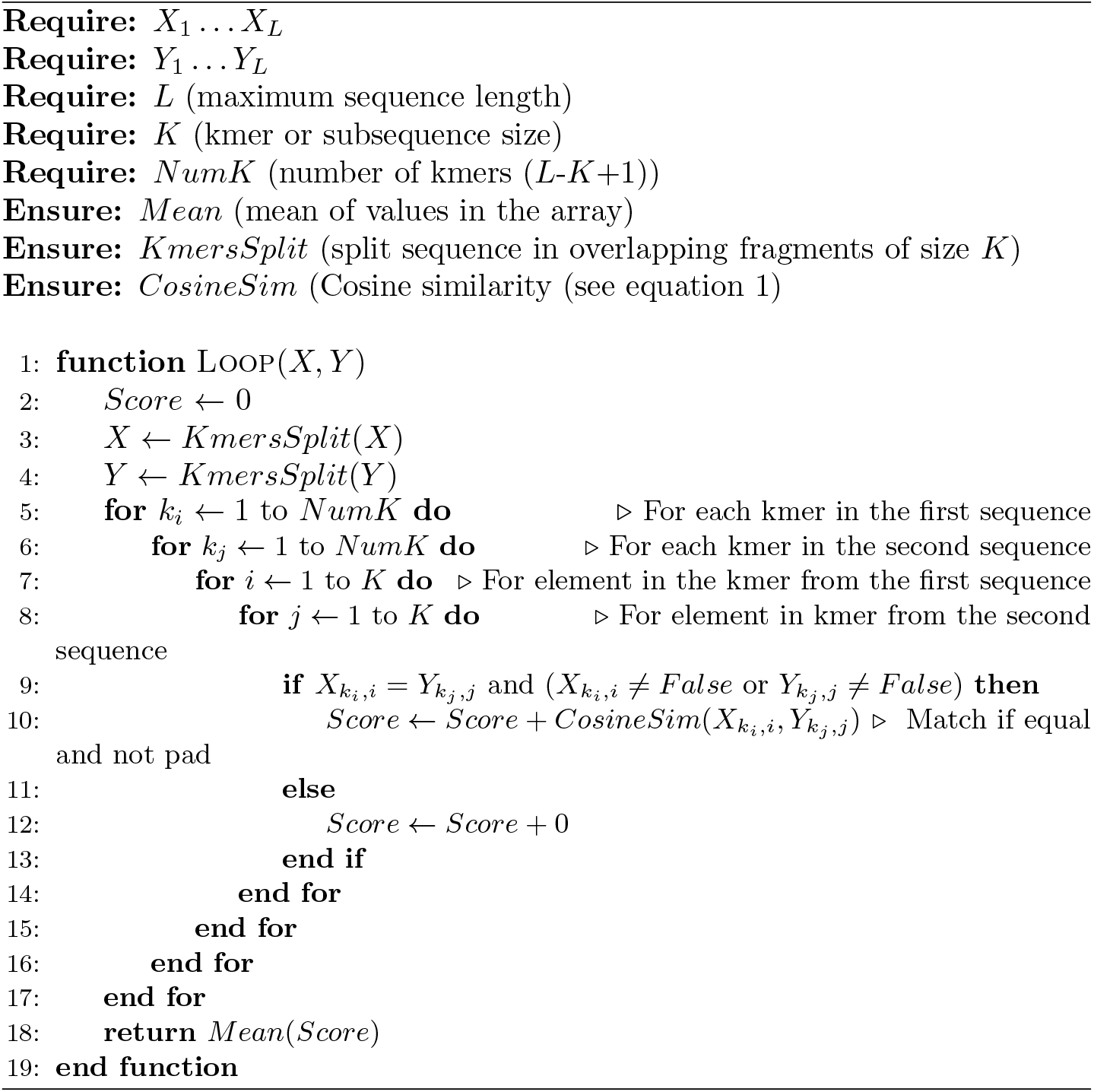

### 3.3 Positional weights

The positional weights are estimated from the previously constructed cosine similarity matrix, without performing the average score among the 2 sequences (see Algorithm 2). Accordingly, we keep track of the pairwise cosine similarity matrix among each element of the compared sequences, building a matrix of size N x N x L x L (see notation at Table 1). Two options are available, if the number of neighbours is set to 1, a column-wise mean is computed to determine the similarity of the current evaluated position against the other elements in the same position (column), excluding itself. Whereas if the number of neighbours is set to 3, then, in addition to the column of the currently evaluated residue, we also take into account the similarity to the columns in the -1 and +1 index positions.

**Table 1:** Notation table

| Name | Description |
| --- | --- |
| $N$ | Number of sequences in the dataset |
| $L$ | Length of the longest sequence in the dataset |
| $K$ | Kmer or subsequence size |
| $NumK$ | Number of kmers of size $K$ in a sequence of size $L$ |
| $D$ | Feature dimensionality size |

## 4 Results

The methods have been tested on different settings for providing an estimation on the required computational time

## 5 Discussion and conclusion

This package has been built due to the lack of the availability of simple packages implementing the cosine similarity among vector encoded sequences (X, Y ∈ R^3^) in python. The method has been able to withstand circa up to 15000 sequences with maximum length 8 and encoded in vectors of size 21 in a time of 24 minutes 26 seconds (see Table 2). Depending on the application, the number of sequences might be traded by longer sequences. The package is expected to expand with new features and improve the memory and time computational requirements in the near future.

**Table 2:** Orientation-table on computational time requirements to simultaneously calculate the 5 similarity measures explained in Section 3. Experiments set up on a Intel(R) Xeon(R) Gold 6136 CPU @ 3.00GHz processor.

| N | L | D | Computational time |
| --- | --- | --- | --- |
| 1000 | 8 | 21 | 9 s |
| 1000 | 50 | 21 | 10 min 25 s |
| 3000 | 40 | 21 | 38 min 25 s |
| 5000 | 8 | 21 | 2min 58 s |
| 15000 | 8 | 21 | 24min 26 s |

## 6 Availability

Examples and code are available at https://github.com/LysSanzMoreta/Dromi

## 7 Acknowledments

Thanks to Flemming Morsch for reviewing and suggesting modifications to the manuscript.

